# Genome-wide identification of heat shock proteins (HSP70) in the Cuban painted snail *Polymita picta*: evolutionary constraints face climate change challenges

**DOI:** 10.64898/2026.01.04.697401

**Authors:** Mario Juan Gordillo-Pérez, Bernardo Reyes-Tur, Zeyuan Chen, Julia Sigwart, Carola Greve, Martijn Heleven, Karen Smeets, Natalie Beenaerts

**Affiliations:** Hasselt University, Centre for Environmental Sciences, Belgium; University of Oriente, Department of Biology and Geography, Cuba; Senckenberg Research Institute and Museum of Natural History Frankfurt, Germany

## Abstract

Climate change poses a critical threat for biodiversity. Among ectothermal animals, terrestrial molluscs are considered highly vulnerable to temperature increase and drought due to their low vagility and specialized ecological requirements. In response to thermal stress, heat shock proteins (HSP70) play a crucial role in the response syndrome and phenotypic plasticity and mediate the adaptation to fluctuating environments. Here, we performed the first genome-wide identification and characterization of the HSP70 family in a Neotropical terrestrial gastropod of high conservation concern: the Cuban painted snail *Polymita picta*. Using comprehensive bioinformatic approaches, we identified HSP70 paralogs, analysed their structural features and assessed sequence variation, highlighting their potential relevance for thermoresistance. Our findings reveal an HSP70 repertoire of ten diversified sequences with signatures of functional specialization and possible adaptive variation linked to environmental temperature regimes. Using novel molecular resources, this pioneering analysis establishes a baseline for early warning systems, integrating stress-related biomarkers into conservation management and climate change adaptation strategies for threatened terrestrial molluscs.

## INTRODUCTION

Ectothermic organisms, such as terrestrial molluscs, are extremely susceptible to fluctuations in temperature and humidity due to lacking acclimation capacity and a limited increase in physiological variance (Morley et al., 2019; Miyahira et al., 2022; Staikou et al., 2024; Noble et al., 2025). Their low vagility, or dispersal capability, poses a challenge for movement and colonization of new habitats with favourable environmental conditions (Kramarenko, 2014). Gastropods are currently exposed to environmental pressures primarily related to human activities, which force them to adapt or acclimate to rapidly changing conditions (Nicolai and Ansart, 2017). However, acclimation does not necessarily result in complete compensation in response to environmental changes (Huey et al., 1999). Often, increases in physiological rates are only partially compensated for (Havird et al., 2020). Disruption of cellular or organismal homeostasis caused by internal or external factors is defined as physiological or primary biological stress (Kultz, 2020). This latter encompasses three major dimensions: intrinsic developmental stress, environmental stress, and aging. The environmental stress arises when external conditions exceed critical thresholds, posing a threat to homeostatic balance (Hayflick, 2007). Within this framework, external factors are termed “stressors,” while the internal reaction they provoke is referred to as “stress” or the “stress response syndrome” (Van Straalen, 2003).

In response to stress, physiological acclimation is mediated by endocrine and epigenetic processes, and relies on a highly evolutionarily conserved stress proteasome that modifies underlying cellular mechanisms to maintain physiological performance (Taff & Vitousek, 2016; Kultz, 2020). It is that populations experiencing high but predictable environmental variability will evolve acclimation, particularly when the fitness costs of plasticity are low (Chevin & Hoffmann, 2017). This capacity has also been associated with various bioecological traits such as habitat type, latitude, body size, and life-history stage (Rohr et al., 2018; Morley et al., 2019).

In general, habitat loss, meteorological phenomena, drought and high temperatures constitute the main threats for terrestrial gastropods affecting species survival, fertility and longevity. Gastropods have evolved several adaptations guaranteeing survival and reproduction in dry and hot environments (Astor et al., 2017; Nicolai and Ansart, 2017). These adaptations involve thermodynamic, physiological, cellular and molecular processes (Schweizer et al., 2019). Aestivation, the induction of chaperones, changes in tissue and cell structure, as well as modifications in mucus production and pigmentation, demonstrate the interrelationship of functions across different levels of biological organization in response to thermal stress and dehydration (Scheil et al., 2012; Bhagat et al., 2016; Schweizer et al., 2019; Tull et al., 2020). Contrasting with the numerous studies on heat effects on macro-physiology and morphology in terrestrial snails, research focusing on tissue, cells or molecular aspects are scarce (Schweizer et al., 2019).

One of the most studied molecular systems directly related to thermal stress response syndrome is the heat shock protein (HSP) family HSP70. These cellular chaperones are ubiquitous, highly conserved proteins distributed in all organisms, from bacteria to humans (Feder et al., 1996; Wheeler and Wong, 2012). According to its variable molecular weight, HSPs are organized in various subfamilies: small HSPs, HSP40, HSP70, HSP90 and HSP110 (Lindquist and Craig 1988). The HSPs with a molecular weight, MW ∼ 70 kDa comprise a set of functionally related molecules known as HSP68, HSP70, HSC70 (cognate heat shock protein), HSP75 (mortalin or mitochondrial heat shock protein) and HSP78 (BiP: binding immunoglobulin or endoplasmic reticulum heat shock protein); all characterized by a highly conserved N-terminal ATPase domain (Wang et al., 2021). In general, HSP70 functions in buffering against intrinsic and extrinsic stress (Murphy, 2013). The roles of HSP70 are variable and specific functions are related to specific stressors. The HSP70 family promotes protein folding and proteastasis, participates in the degradation of misfolded sequences and intervenes in translocation, and prevents deleterious aggregation (Mayer and Bukau, 2005; Li et al., 2025). Additionally, essential functions in response to environmental stress such as heat, hypoxia, pathogen infection, exposure to organochlorines and heavy metals, physiological stress and immune responses have also been observed (Moseley, 1997; Mosser et al., 1998; Tsan and Gao, 2009; Deane and Woo, 2011; Chen et al., 2018; Shi et al., 2020). In molluscs, HSP70 proteins are localized in cytoplasm, nuclei, endoplasmic reticulum and mitochondrion and each subcellular location is determined by specific motifs (Rosenzweig et al., 2019).

The HSP70 proteins are classified into two distinct groups based on their function and regulation patterns: constitutive expressed heat shock proteins (HSCs), which are expressed in the cells under normal conditions, and inducible heat shock proteins (HSPs), which are expressed under specific stress conditions (Boone and Vijayan, 2002). In general, HSP70 contains three main conserved domains: N-terminal ATPase domain (∼ 40 kDa), also called the nucleotide-binding domain (NBD); it controls the interaction of the chaperone with the client protein. A second substrate-binding domain (SBD) (∼ 18 kDa), binds hydrophobic regions of client proteins during early folding. And lastly, a third C-terminal domain with variable length (∼10 kDa) (Nikolaidis and Nei, 2004; Daugaard et al., 2007; Melikov et al., 2024). Until now, HSP70 in terrestrial molluscs has been predominately studied via Western blot in Mediterranean species (e.g. Di Lellis et al., 2012, 2014; Dieterich et al., 2013, 2015; Staikou et al., 2024). The development of genomic technologies and the recent increasing availability of whole genome sequences of mollusc species, enables us to assess the HSP70 gene family within an evolutionary context and its role in the adaptation to different environments (e.g. Hu et al., 2022; Lu et al., 2024).

In the Neotropical Cuban archipelago, the genus *Polymita* Beck, 1837, includes six endemic tree snail species: *P. muscarum*, *P. versicolor*, *P. venusta*, *P. sulphurosa*, *P. brocheri* and *P. picta* (González-Guillén, 2021) all inhabiting East Cuba to northeast of Sabana-Camaguey (de la Torre, 1950). The “polimitas” are admired worldwide because of their remarkable shell colour polymorphism (Alfonso y Berovides, 1993), arousing the interest of evolutionary biologists, conservationists and collectors. They were recently awarded as Mollusc of the Year 2022 (Mayer-Bömoser et al., 2022). All *Polymita* species are considered threatened (Maceira-Filgueira, 2016) and listed in Appendix I by the Convention on International Trade in Endangered Species (CITES, 2022).

Since its description as *Helix* (*Polymita*) *picta* (Born, 1778), this charismatic species has received growing research interest from different disciplines: taxonomy, behaviour, reproductive anatomy, shell polymorphism and ecology (e.g. Andrews, 1932; de la Torre, 1950; Moreno, 1950; Berovides et al., 1986; Alfonso and Berovides, 1987, 1989a, 1989b; Berovides and Alfonso, 1992; Berovides et al., 1986). Recently, its shell’s luminance was quantified and its potential link to thermoresistance was discussed (Gordillo-Pérez et al., 2025). Simultaneously, the full sequencing of the mitochondrial and nuclear genes, provided new insights into its phylogeny (Lewis et al., 2025; Reyes-Tur et al., 2025).

In a recent study on six Cuban species of land snails found that ecological niche models predicted a loss of favourable habitats for all species by 2050 (Martínez y Hernández 2017). Besides, Mancina et al. (2017), conducted similar research on the six *Polymita* species, reporting habitat decrease in all cases and the vulnerability status of the genus in three hypothetical climate change scenarios for 2050. Despite the effort made in research and conservation of Cuban land snails, there are still no molecular studies investigating the resilience of these animals to climatic variability.

The aim of this paper is to conduct a genome-wide identification of the 70-kDa heat shock protein (HSP70) family in *Polymita picta*, considering its role in thermal stress response and phenotypic plasticity as key mechanisms of climate-change adaptation. As far as we know, this research explores for the first time an HSP family at genome level in a Neotropical terrestrial gastropod.

## MATERIAL AND METHODS

### Genome-wide identification of HSP70 family members in the *Polymita picta* genome

The identification of HSP70 proteins in the *P. picta* genome was based on sequence homology and conserved domain analysis following methodologies carried out in similar studies (Hu et al., 2022; Lu et al., 2024). The genome of *Polymita picta* was obtained from NCBI (https://www.ncbi.nlm.nih.gov/bioproject/PRJNA1250545), and a genome-wide TBLASTN search was performed using all available mollusc HSP sequences from NCBI (1,064 sequences). The resulting TBLASTN hits (78,994) were filtered in R (R Core Team, 2022) to retain sequences with an E-value ≤ e-5.

To verify the presence of conserved protein domains in the candidate sequences, a hidden Markov model (HMM) profile for HSP70 (PF00012) was downloaded from the Pfam protein family database (https://pfam.xfam.org/), and processed using the software HMMER 3.0. The presence of conserved HSP70 domains was further confirmed using the NCBI Conserved Domains Database (http://www.ncbi.nlm.nih.gov/cdd/), with an E-value of <0.0001 and all other parameters set to their default values.

### Full-length sequence extraction and translation from genomic data

The 56 HSP70 protein fragments identified through HMM were re-aligned to the *P. picta* genome assembly (version v4665_2) in NCBI using BLAST+ v2.13.0 (TBLASTN) (Camacho et al., 2009). The corresponding BLAST hits were provided in BED format, containing genomic coordinates for each identified region. The BED file containing the genomic coordinates of the HSP70 hits was used to extract the corresponding DNA sequences from the genome. Each hit provided the start and end positions for sequence extraction. If the sequence was identified on the negative strand of the genome, it was reverse complemented to ensure proper reading frame translation. To identify the best open reading frame (ORF), we tested three possible reading frames for each extracted sequence. This was done by translating the sequence from three different starting positions corresponding to the three potential frames. The number of stop codons (“*”) was counted in each translated sequence. The reading frame with the fewest stop codons was selected as the optimal frame for translation. Sequences with internal stop codons were discarded, and the longest continuous sequence without stop codons was retained for further analysis. After selecting the optimal reading frame for each sequence, the corresponding protein sequence was translated (Supplementary Data 1: https://doi.org/10.5281/zenodo.16744586). These procedures were carried out in R software (R Core Team, 2022). We manually checked the R extraction using Integrative Genomic Viewer (IGV) ver. 2.19.3. (Thorvaldsdóttir et al., 2013). Each translated sequence was then annotated with its genomic coordinates (scaffold, start, end, and strand direction).

### Physicochemical profiling of *Polymita picta* HSP70s

The physicochemical properties, including molecular weight (MW), number of amino acids, theoretical isoelectric point (pI), grand average of hydropathy (GRAVY), and instability index (II), were analysed using the ExPASy ProtParam tool. The signal peptide prediction i n *P. picta* HSP70s (*Pp*HSP70) was carried out through the online tool Signal P 5.0 (http://services.healthtech.dtu.dk/service.php?SignalP-5.0. The conserved motifs in *Pp*HSP70 were analysed using the Multiple EM for Motif Elicitation online tool (http://meme-suite.org/meme/tools/meme).

### Motifs analysis in *Polymita picta* HSP70s and comparative sequences

A total of 34 protein sequences were analysed for conserved motifs, including the 10 HSP70 paralogs from *P. picta* and additionally HSP70 sequences from the molluscan species *Biomphalaria glabrata*, *Ostrea edulis*, *Magallana gigas*, *Mya arenaria*, *Aplysia californica*, *Dreissena polymorpha* a n d *Crassostrea virginica* and one HSP70 from *Drosophila melanogaster*. For helping identifying subfamilies motifs, sequences annotated as HSP70 cytosolic, HSC70, HSP75 and HSP78 were included (Supplementary Data 2: https://doi.org/10.5281/zenodo.16744775). We selected the aforementioned species to enable orthologous-based comparisons of HSP genes with those of *Polymita*, prioritizing taxa with available reference sequences and leveraging the high evolutionary conservation of HSPs (Wheeler and Wong, 2012).

### Homology and gene duplication

Two criteria were considered to identify gene duplications: (i) sequence similarity greater than 70% and (ii) the shorter aligned sequence covering at least 70% of the longer one (Gu et al., 2002; Yang et al., 2008; Lu et al., 2024). The sequences were aligned using MAFFT v7 (https://mafft.cbrc.jp/alignment/server/index.html) and loaded into R using the Biostrings package. Metadata including sequence names, scaffolds, and strand orientation were parsed from the FASTA headers using regular expressions and stored in a structured data frame. A pairwise comparison between all sequences was conducted to compute percentage identity. A custom R function compared aligned amino acids for each pair and calculated identity as the proportion of matching positions over the total alignment length (R Core Team, 2022). In all pairs selection pressure rates for duplicated genes were estimated using Ka/Ks_Calculator 3.0.

### Phylogenetic analysis

For the phylogenetic analysis, the sequences of *P. picta* HSP70 that had been identified were re-aligned to the same set of orthologous used for motif identification, ensuring consistency across comparative analyses. The major heat shock 70 kDa protein from the model species *Drosophila melanogaster* was used as an out-group. The LG+G4 amino acid substitution model was selected for phylogenetics through the ModelFinder function (-m MFP) (Kalyaanamoorthy et al., 2017) followed by tree inference using IQ-TREE v2.3.6 (Nguyen et al., 2015; Minh et al., 2020). Branch support was estimated through ultrafast bootstraps (Hoang et al., 2018) and Shimodaira-Hasegawa like approximate likelihood ratio test (SH∼aLRT) (Guindon et al., 2010). In both algorithms 10 000 replications were performed. Nodes with an ultrafast bootstrap value ≥ 95 and a likelihood ratio ≥80 were considered as well as supported (Cruz-Laufer et al., 2025). Finally, phylogenetic trees were visualized using FigTree v1.4.4.

## RESULTS

### Genome-wide identification and physicochemical profiling of heat shock proteins 70 kDa (HSP70) in *Polymita picta*

In this study, 56 partial sequences of HSP70 were identified across the genome of *P. picta* using a TBLASTN search with 1,064 mollusc sequences from NCBI. After screening using HMM and CDD-NCBI, 22 sequences (39.2%) showed significant similarity to a query containing the ATPase-like domain of the ASKHA superfamily (Acetate and Sugar Kinases/Hsc70/Actin), while the remaining sequences (60.8%) were associated with the PTZ00009 domain, which is also annotated for heat shock 70 kDa proteins.

Heat shock proteins were identified at 10 unique positions in the genome and were named consecutively *Pp*HSP70_1 to *Pp*HSP70_10. Information on the identifier is followed by the scaffold where it is located, the start and end coordinates, and the strand, e.g., *Pp*H SP 70 _ 1 _ptg 000811 l_ 1: 167165 - 168904 (-) (Supplementary Data 3: https://doi.org/10.5281/zenodo.16744894).

The ten HSP70 protein sequences identified were distributed across five scaffolds (Fig. 1). *Pp*HSP70_1 to *Pp*HSP70_6 were closely located on both strands of ptg000811l_1 (only *Pp*HSP70_5 and *P*pHSP70_6 on the positive strand), while the remaining four sequences were distributed across the positive strand of scaffolds ptg004791l_1, ptg010661l_1, ptg018870l_1, and ptg018978l_1.

**Figure 1.**
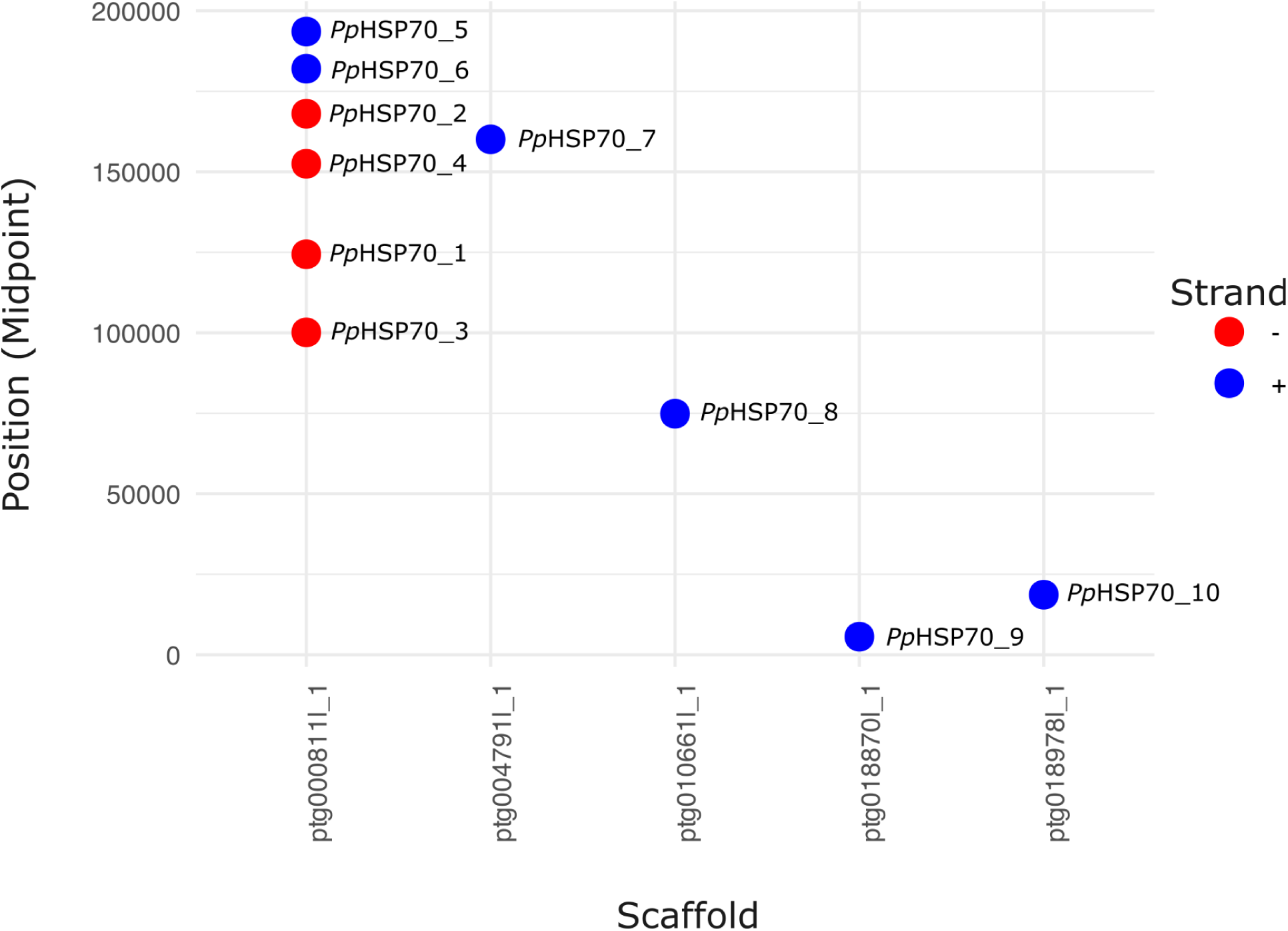
Scaffold distribution of *Polymita picta* heat shock proteins 70 kDa (HSP70).

The 10 different identified HSP70 proteins consist of amino acid sequences ranging from 635 to 672, with an average of approximately 641 residues (Table 1). The molecular weights varied between 69,305.92 Da and 73,549.12 Da, with an average of 69,725.06 Da. The isoelectric points (pI) were consistently low, ranging from 5.58 to 5.80 (mean pI = 5.67), indicating a generally acidic nature. Additionally, all proteins exhibited negative GRAVY values (–0.47 to –0.38; mean = –0.44), suggesting overall hydrophilicity. The instability indices for all sequences were below 40, ranging from 28.23 to 30.69 (mean = 29.24), classifying them as stable proteins. The aliphatic index values ranged from 79.43 to 84.32 (mean = 81.15), supporting moderate to high predicted thermostability (Table 1).

**Table 1.** Physico-chemical parameters of *Polymita picta* heat shock protein 70 kDa.

| Sequence | Number of Amino Acids | Molecular Weight | Isoelectric Point | GRAVY | Instability Index | Aliphatic Index |
| --- | --- | --- | --- | --- | --- | --- |
| ***PpHSP70_1*** | 637 | 69536.36 | 5.66 | -0.44 | 28.81 | 81.46 |
| ***PpHSP70_2*** | 637 | 69550.26 | 5.58 | -0.46 | 30.69 | 81 |
| ***PpHSP70_3*** | 645 | 70355.17 | 5.59 | -0.45 | 29.52 | 80.76 |
| ***PpHSP70_4*** | 637 | 69607.35 | 5.66 | -0.47 | 29.84 | 80.55 |
| ***PpHSP70_5*** | 637 | 69580.28 | 5.58 | -0.46 | 30.2 | 80.55 |
| ***PpHSP70_6*** | 637 | 69607.35 | 5.66 | -0.47 | 29.84 | 80.55 |
| ***PpHSP70_7*** | 672 | 73549.12 | 5.8 | -0.38 | 29.44 | 84.32 |
| ***PpHSP70_8*** | 635 | 69305.92 | 5.76 | -0.47 | 28.23 | 79.43 |
| ***PpHSP70_9*** | 641 | 69925.64 | 5.69 | -0.44 | 28.44 | 80.34 |
| ***PpHSP70_10*** | 635 | 69317.22 | 5.77 | -0.42 | 30.34 | 83.56 |

Al l *P. picta* HSP70 proteins appear to be primarily localized in the cytoplasm, with prediction scores ranging from 0.47 to 0.78 (mean = 0.73) (Supplementary Data 4: https://doi.org/10.5281/zenodo.16745227). However, two proteins, *Pp*HSP70_3 and *Pp*HSP70_8, showed nuclear localization signals (NLS), implying potential dual localization in both the cytoplasm and nucleus. Additionally, *Pp*HSP70_7 displayed a peroxisomal targeting signal, with a relatively lower cytoplasmic probability (0.47), but higher probabilities for other compartments such as the endoplasmic reticulum (0.52), Golgi apparatus (0.47), and lysosome/vacuole (0.46). Probabilities for mitochondrial or cell membrane localization were generally low across all proteins (mean < 0.34).

Signal peptide prediction using SignalP 5.0 indicated that none of the *Pp*HSP70 proteins possess a classical N-terminal signal peptide for secretion via the Sec/SPI pathway (General secretion pathway / Signal Peptidase I). All sequences were classified as “OTHER” with high probability (average OTHER probability = 0.9975 ± 0.0003) (Supplementary Data 5: https://doi.org/10.5281/zenodo.16745405). The SP (Sec/SPI) scores were consistently low across all sequences (average SP probability = 0.0025 ± 0.0003), suggesting that these proteins are not secreted through the classical secretory pathway.

### Motif Analysis of HSP70 Family in *Polymita picta* and Comparative Sequences

A total of 21 motifs were identified across the dataset, revealing distinct differences in motif architecture that correlate with subcellular localization and functional HSP70 subfamilies. Notably, mitochondrial HSP70s (HSP75) shared a unique motif at the N-terminal region (motif 21), which was absent from all other sequences. Both HSP75 and HSP78 (endoplasmic reticulum) displayed an additional motif in the IIA Actin-like ATPase subdomain of NBD, which was not found in cytosolic HSP70s, including those identified in *P. picta*. The three HSP70 cognate proteins (HSC70) lacked a conserved motif (motif 15) in the SBD, which was consistently present in all inducible HSP70s, including all *P. picta* paralogs and the *D. melanogaster* inducible HSP70.

Minor variations were detected in the height and sequence of two short motifs (∼8 amino acids), located near the IA Actin-like ATPase subdomain of NBD and in the central region of the SBD. These changes correspond to single amino acid polymorphisms (SAPs). Sequence alignment (Supplementary Data 6: https://doi.org/10.5281/zenodo.16746341), confirmed that these SAPs are scattered and mostly conservative. For example, in the C-terminal region, just before the EEVD sequence (see motif 16 in Fig. 2), an insertion of four or five residues was detected between positions 638-642 in PpHSP70_7 (SGSSR), *Pp*HSP70_8 (AHSQ), *Pp*HSP70_9 (AGSQ), and *Pp*HSP70_10 (TGGHS). Additionally, a residue deletion at position 635 is observed in *Pp*HSP70_10, and a second deletion at position 627 appears in *Pp*HSP70_7, *Pp*HSP70_8, and *Pp*HSP70_10. The N-terminal region displayed slight variability in the residues following the initiating methionine: *Pp*HSP70_1 starts with MA, *Pp*HSP70_2 with MR, *Pp*HSP70_5 with MT, and *Pp*HSP70_6 with MS. Notably, *Pp*HSP70_10 lacks the first five amino acids.

**Figure 2.**
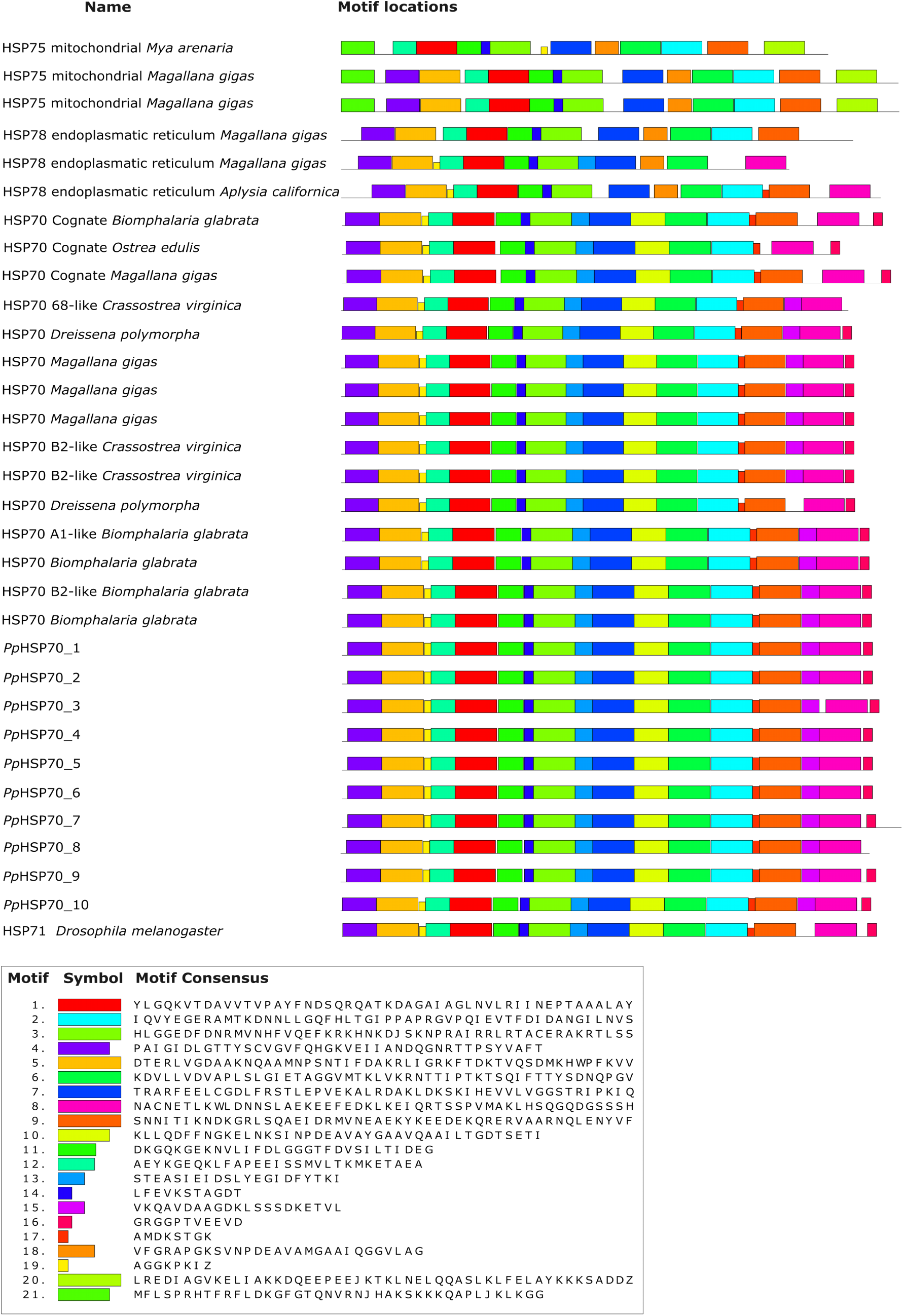
Distribution of conserved motifs of different subfamilies of heat shock proteins 70 kDa (HSP70) in *Polymita picta* and comparative sequences.

### Sequence Identity and Evolutionary Constraints

We computed pairwise amino acid identity percentages to assess the degree of similarity among the identified HSP70 paralogous proteins in *P. picta*. As shown (Fig. 3), proteins located within the same scaffold (e.g., ptg000811l_1) generally exhibit very high identity (>97%). In contrast, lower identity values were observed for sequences located in different scaffolds. The minimum identity, 83.26 %, was found between *Pp*HSP70_7 and *Pp*HSP70_10.

**Figure 3.**
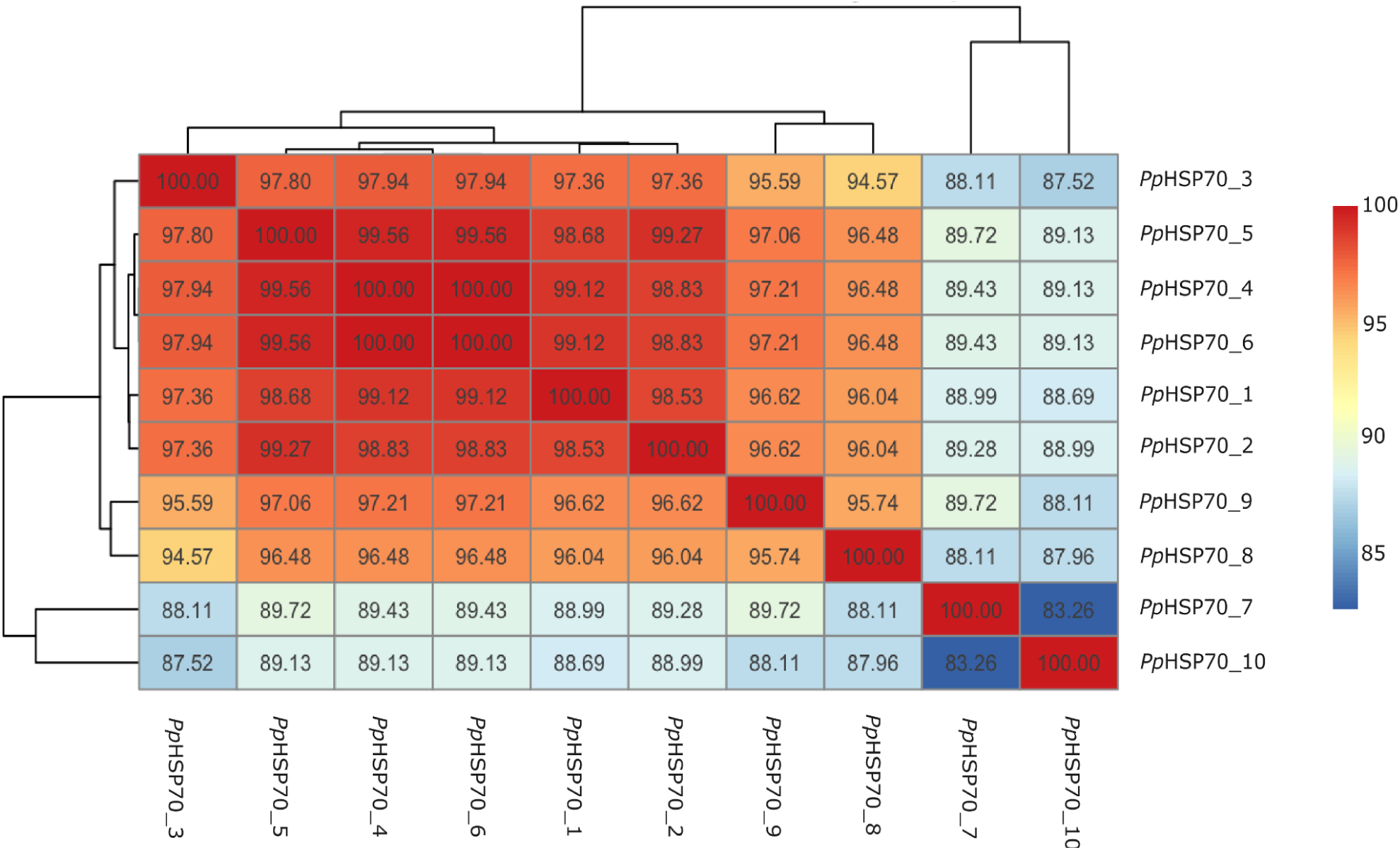
Matrix of identical positions (pID) percentages between *Polymita picta* heat shock protein 70 kDa (HSP70) sequences.

The Ka/Ks ratios for the various HSP70 sequence pairs were calculated to assess selective pressure and potential evolutionary divergence. The results, presented in (Supplementary Data 7: https://doi.org/10.5281/zenodo.16751419), include the comparison of 45 pairwise combinations of HSP70 sequences in *P. picta*. As a general trend, all calculated Ka/Ks ratios are below 1. For example, the sequence pair *Pp*HSP70_1 vs *Pp*HSP70_10 exhibited the lowest ratio of 0.0127. In contrast, the pairs *Pp*HSP70_3 vs *Pp*HSP70_4 and *Pp*HSP70_3 vs *Pp*HSP70_6 both exhibited the highest Ka/Ks ratio of 0.1642. Notably, the pair *Pp*HSP70_4 vs *Pp*HSP70_6 returned a NaN value due to the sequences being identical but located at different loci.

### Phylogenetic analysis of *Polymita picta* HSP70 and comparative sequences

The phylogenetic analysis revealed the presence of two well defined major clades (Fig. 4). The first clade grouped all *P. picta* sequences together with four cytosolic HSP70 sequences from the pulmonate *B. glabrata*, with strong statistical support (SH-aLRT/UFBOOT = 100/100). Within this clade, two main branches are identified: one branch contains exclusively the four *B. glabrata* sequences, supported by very high values (SH-aLRT/UFBOOT = 100/100 and 98.8/100) and the other branch grouping the ten *Pp*HSP70 sequences. The internal relationships among the *P. picta* sequences display heterogeneous support. In particular, the placement of PpHSP70_10 relative to the other paralogs exhibited low resolution. Similarly, the lowest bootstrap values were observed at the nodes grouping *Pp*HSP70_1, *Pp*HSP70_3, *Pp*HSP70_4, and *Pp*HSP70_6. By contrast, the remaining nodes within the paralogs clade showed better resolved relationships. The second major clade comprised sequences from all other molluscan species corresponding to HSC, HSP75, and HSP78 also with high support (SH-aLRT/UFBOOT = 99.4/100). Within this clade, a secondary branch grouped other cytosolic HSP70 sequences, although their internal relationships remained unresolved.

**Figure 4.**
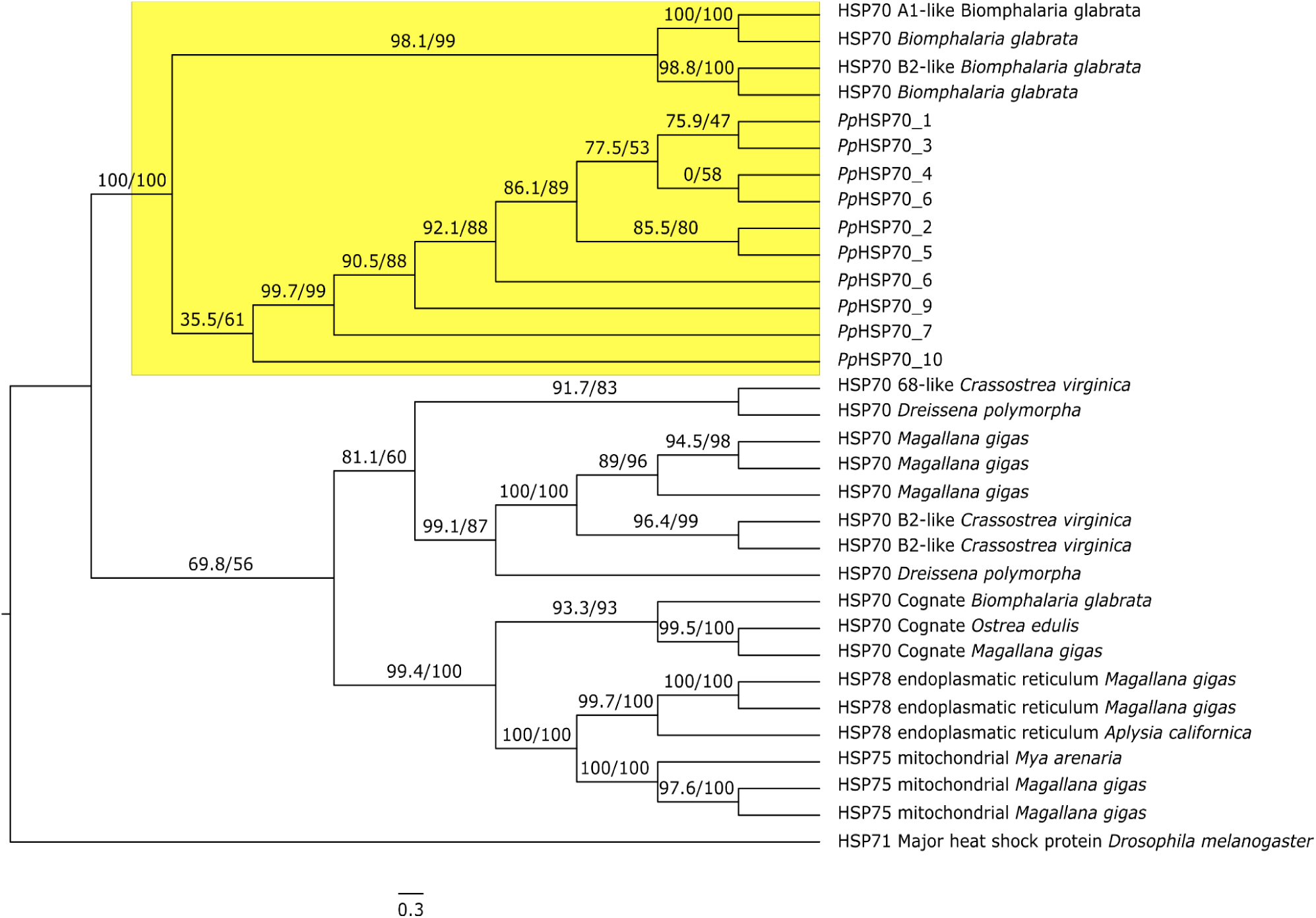
Phylogenetic analysis of *Polymita picta* heat shock protein 70 kDa (HSP70) amino acid sequences and orthologous. The node grouping *P. picta* sequences is highlighted in yellow.

## DISCUSSION

To ultimately provide policy makers with sound advice on the protection of vulnerable fauna in a changing climate, more specifically in the sub-tropics, we focused on the HSP70 protein family, given its promising roles in coping with the physical challenges caused by climate change, of the Cuban painted snail *Polymita picta*. Therefore, we performed a genome-wide identification of the chaperones and analysed its molecular characteristics, variation and evolution.

### The genome-wide identification of HSP70s in *Polymita picta* and its potential relationship with thermoresistance

The heat shock proteins of 70 kDa (HSP70) is a well characterized protein family playing a crucial role in the cells’ housekeeping and stress response, from bacteria to mammals (Karlin and Brocchieri, 1998; Clerico et al., 2019; Rosenzweig et al., 2019). However, until now related studies in molluscs mainly focused on species of economic interest being a food source or a threat as an invasive species (Zhang et al., 2012; Cheng et al., 2016; Hu et al., 2019, 2022; Lu et al., 2024). Unfortunately, studies addressing the HSP70s in land snails from a genomic angle are totally lacking. This evidently complicates understanding the evolution and diversity of these essential molecules.

Besides many new research opportunities, the recent *P. picta* full genome sequencing (Reyes-Tur et al., 2025), facilitated this first HSP70 screening. The genome assembly contained 1.85 Gbp with 22628 contigs and a contig N50 of 124.2 Kbp. Moreover, the genome completeness corresponds to 88.4 % and the high heterozygosity, 2.41 % are as expected (Vendrami et al., 2020; Bean et al., 2022; Gundappa et al., 2022; Reyes-Tur et al., 2025). Therefore, we interpreted our results at scaffold level and conservatively (to avoid identifying alleles as duplications). We identified 10 different genes coding for full length heat shock proteins The number of HSP70 variants retrieved, is comparable with those found in several fish species; or even the sea cucumber *Apostichopus japonicus* (Xu et al., 2018; Ma et al., 2020; Gao et al., 2018, 2022; Zheng et al., 2023). Strikingly, in bivalve species the number of observed HSP70s proteins, including several tandem duplicated sequences, is manyfold e.g. 65 genes in the clam *Telligarga granosa* (Kim et al., 2024) or 113 variants in the pacific oyster *M. gigas* (Lu et al., 2024).

The 10 HSP70 identified in this study were distributed in five different scaffolds. Four sequences, *Pp*HSP70_7 to *Pp*HSP70_10 were located in four of the five different scaffolds, which suggest they represent putative dispersed duplications, being majorly a product of transposition (Wang, 2011). The other sequences, from *Pp*HSP70_1 to *Pp*HSP70_6 are located in the same scaffold in each other’s vicinity (< 10 bp apart); four of these in the negative strand and two in the positive one. Most likely, this study included tandem duplications (Reams et al., 2015), a common phenomenon well described in mollusc genomes (Zheng et al., 2022; Lu et al., 2024). Consequently, providing them with functional variation, relevant for adapting to different and/or changing environments, coping with diseases and potentially useful in conservation management (Hawkins et al., 2024; Sutherland et al., 2024, 2025).

Overall, *P. picta* HSP70 proteins show high conservation in size, with an average of 641 residues, low isoelectric points indicating their acidic nature and negative GRAVY values, which suggest their hydrophilic character. Therefore, all molecules can be classified as stable proteins. Additionally, due to their aliphatic indexes we consider them to have a moderately to high thermostability. These physico-chemical characteristics are typical of the HSP70 family (Dayrit et al., 2023; Singh et al., 2025).

The 10 identified sequences are most likely located in the cytoplasm, however, two proteins, *Pp*HSP70_3 and *Pp*HSP70_8 also showed signals corresponding to a nuclear position suggesting a potential dual function both in cytoplasm and nucleus. Moreover, one sequence, *Pp*HSP70_7 may have a broader subcellular distribution, showing target signals for peroxisome, endoplasmic reticulum, Golgi apparatus and lysosome/vacuole. This possibly reflects a specialized or a distinct stress-related function from the other members (Daugaard et al., 2007). *Polymita picta*’s HSP70 proteins therefore can be considered as predominantly cytoplasmic chaperones, with potential minor roles in other organelles for a subset of sequences (Peyretaillade et al., 1998; Kaneko, 2022; Hasnain et al., 2022; Yadav et al., 2019). Importantly, our results confirm that none of the ten sequences is secreted through the classical secretory pathway (Sec/SPI), supporting our conclusion that all *Pp*HSP70 in *P. picta* are primarily cytoplasmic chaperones.

### Motifs analysis of HSP70 family in *Polymita picta* and comparative sequences

The identified motifs correspond to canonical HSP70 domains, supporting functional conservation across the family despite gene duplications. The unique motifs in organellar subfamilies and the absence of specific ones for HSC70s highlight the structural divergence between cytosolic and non-cytosolic members of the HSP70 family. For the case of this chaperone family, major differences are observed at C-terminal. The three HSP70 cognate proteins (HSC70) included in our analysis lacked a conserved motif (motif 15) in the SBD, consistently present in all inducible HSP70s, including all *P. picta* paralogs and the *D. melanogaster* inducible HSP70, what suggests functional divergence between constitutive and inducible variants. Some identified changes among *P. picta* paralogous are based on single amino acid polymorphisms (SAPs) that do not disrupt motif presence but may indicate allelic or functional divergence. With the exception of *Pp*HSP70_8, which lacks the motif 16 and exhibits some differences, potentially related to the exon-intron structure of the C-terminal region (see distance between motifs 8 and 15 in *Pp*HSP70_3), the motif patterns are conserved. This suggests that the identified chaperons in *P. picta* have unique biological functions that vary discretely (Xu et al., 2018; Zhang et al., 2013; Huang et al., 2018). Notably, *Pp*HSP70_10 lacks the first five amino acids, potentially indicating an alternative start codon but the sequence possesses the full first identified motif on N-terminal (motif 4).

Despite these minor substitutions, no major insertions or deletions were detected, and all sequences retained the expected domain architecture and length consistent with functional HSP70 proteins. The preservation of key domains such as the nucleotide-binding domain, substrate-binding domain, and the C-terminal EEVD motif suggests that all sequences maintain their chaperone function. Taken together, the high degree of sequence and motif conservation, combined with modest SAPs, supports the hypothesis that the HSP70 sequences in *P. picta* may represent recent gene duplications or allelic variants within a smaller number of loci in comparison to marine molluscs species. Terrestrial ectotherms are generally less able to acclimatize and exhibit smaller increases in physiological rates than aquatic ones (Noble et al., 2025). In any case, further genomic, transcriptomics, and expression analyses are necessary to elucidate the functional relevance of the observed polymorphisms. Motif-based comparisons between *Polymita* HSPs sequences and orthologs provide preliminary context but should be interpreted with caution. Our analyses were necessarily constrained by the availability and quality of reference sequences and functional inferences from motif presence/absence remain indirect. Accordingly, we use the existing experimental literature to contextualize putative roles and to guide hypotheses about protein function. More functionally informative comparisons - specially with orthologs from molluscan lineages most closely related to *Polymita* (e.g. terrestrial gastropods) - will depend on the continued expansion of high quality genomic and transcriptomics resources for these taxa.

### Sequence Identity and Evolutionary Constraint

The amino acid sequences of *Pp*HSP70_1 to *Pp*HSP70_6 found in the same scaffold demonstrate high levels of homology (>97%). This suggests either recent gene duplications, as previously proposed, or strong functional constraints. Despite the possible presence of allelic variants among these *P. picta* sequences, tandem duplication of HSP70 genes has been reported in several molluscs species. For example HSP70 genes are distributed across all 19 chromosomes of *M. mercenaria*, and the HSPa12 gene appears to be duplicated 39 times on chromosome 7 (Hu et al., 2022). A similar high-density collinear gene block has been observed in the chromosome 14 of *C. sinensis* (Wei et al., 2020), as well as in other bivalves belonging to the subclass Pteriomorpha (marine species) and the order Veneroida (marine and fresh water species) (Zhang et al., 2012; Cheng et al., 2016; Powell et al., 2018; Peng et al., 2020; Gao et al., 2022). Lower identity values were found in the sequences from *Pp*HSP70_7 to *Pp*HSP70_10, but still presenting more than 83 % of homology, suggesting older duplication events followed by varying degrees of sequence divergence. It is understood that duplications can exhibit homology values ranging from 70 to 100 % in the case of HSP70 proteins (Lu et al., 2024).

All Ka/Ks ratios between *Pp*HSP70 pairs are below 0.16, indicating strong purifying selection. The duplicated genes typically remain under purifying selection during the early stages of evolution (Kondrashov et al., 2002), to ensure normal cellular function, while those that respond properly to stress tend to be amplified (Evgen’ev et al., 2014; Metzger et al., 2016). It is well known, gene duplication provides material for evolutionary novelty and tandem duplications is recognized as an important driving force in this process (Roth et al., 2007; Chen et al., 2013; Long et al., 2013; Singh et al., 2025). Generally, variability in HSP70 gene copy number is assumed to reflect regulatory differences, particularly in relation to the factor temperature. Furthermore, genomic variability resulting from gene duplications has been reported to directly influence physiological responses to environmental fluctuations (Zhang et al., 2012). Based on the observed homology among *P. picta* HSP70 sequences and the selective pressure ratios, it appears that the gene duplication events in this species are recent and still subject to purifying selection

Our preliminary genomic evidence suggests that *P. picta* may harbor an expanded HSP70 gene family relative to molluscs that routinely experience broad thermal regimes (e.g. intertidal oysters), yet *Polymita* inhabits comparatively thermally stable terrestrial microhabitats. To elucidate how the architecture of the *Pp*HSP70 family relates to the species’s life history, it is necessary to consider both, the evolutionary trajectory of *Polymita* in prehistoric Eastern Cuba and the evolution of its habitats. Historically, the archipelago has experienced profound paleoclimatic fluctuations since early Miocene, when the origin of its modern fauna is estimated (MacPhee & Iturralde-Vinent, 2005). Episodes of tectonic uplift, Pleistocene sea-level oscillations and recurrent drought-humid cycles recorded in speleothem and lacustrin archives (Fensterer et al., 2013; Peros et al., 2015, 2017) likely shaped population connectivity, niche distributions and thermotolerance in ancestral *Polymita* lineages. It is therefore expected that the basal architecture of the HSP70 family was modeled under these long term selective regimes. Moreover, the evolution of thermal stress genes has also been linked to ecological drivers beyond temperature per se (Staikou et al., 2024). Future functional studies that profile differential expression of individual paralogs across environmental gradients will be crucial to disentangle the relative contribution of ancient versus ongoing selective pressures to *Polymita*’s HSP70 complement.

### Conservation perspectives

In terms of conservation, over the past centuries, Cuba has undergone dramatic habitat transformation, including the loss of ∼ 85 % of native forests within seven decades (del Risco, 1995) and persistent deforestation-driven fragmentation of habitats. Although recent reports indicate a slight recovery of forest cover, ∼ 32.2 % (ONEI, 2025), climatic stressors are intensifying: annual mean temperatures are rising, heat extremes are breaking historical records and rainy season shows increased thermal variability (Centella et al., 1999; Febles, 2009; Fonseca-Rivero et al., 2024; Harbott et al., 2023; Campbell, 2023).

Moreover, Cuba’s protected-area system covers only 15.94% of the national territory (ONEI, 2025), in a region that forms part of a biodiversity hotspot. Protected areas do not always encompass the full extent of species’ distributions (Mancina and Cruz, 2017), and interactions with surrounding ecosystems remain poorly investigated. In addition to this, the country has experienced deep political and economic crises in recent decades, resulting in a shortage of vital resources and hunger, particularly in rural areas. In response, the use of firewood for cooking and charcoal production as surviving strategies, lack of and deficiencies in management plans have become a direct threat to the forest ecosystems (Maal-Bared, 2006; Alonso and Clark, 2015).

In an adaptive context, evidence of an expanded HSP70 chaperone family in *Polymita* does not, by itself, imply resilience to climate change or heat stress. Gene duplication is common but only rarely yields clear adaptive gains and increase in copy number can be neutral or even deleterious (Qian and Zhang, 2014). Classic experiments with *Drosophila* further show that adding HSP70 copies can improve survival under severe heat stress yet reduce performance under benign or moderately stressful conditions-underscoring that “more copies” does not straightforwardly translate into greater thermal tolerance- (Feder et al., 1996; Roberts et al., 2000). More broadly, thermal tolerance itself limited plasticity across ectotherms, cautioning against equating gene-family size with climate resilience (Gunderson & Stillman, 2015).

The finds presented in this research regarding *P. picta* HSP70, are the result of a genome-wide identification in one individual used for the genome sequencing (Reyes-Tur at al., 2025); so, intra and interspecific variations in the HSP70 system on the genus and the study of other stress related molecules should be taken into account for a better understanding of the thermoresistance molecular mechanisms. We understand that more in-depth experimental research is needed to improve our understanding of *Polymita*’s resilience to global warming in the Anthropocene, and more specifically in the context of Cuba. The early detection of “more thermoresistant” haplotypes could constitute a strategy for genetic conservation, population reinforcement and re-introduction.

